# THE STRIATAL HETERODIMERS A_2A_R-D_2_Rs ARE LINKED TO ERGOGENIC EFFECTS OF CAFFEINE

**DOI:** 10.1101/2023.05.24.541930

**Authors:** Ana Cristina de Bem Alves, Naiara de Souza Santos, Ana Paula Tavares, Gabriela Panatta, Aderbal S Aguiar

**Author notes:** **Corresponding author** Ana Cristina de Bem Alves, PhD. Department of Health Sciences, Federal University of Santa Catarina (UFSC), Ararangua, 88905-120, SC, Brazil. Cel.

## Abstract

Adenosine 2A receptors (A_2A_Rs) are abundant in the striatum, and they are found co-localized with dopamine D2 receptors (D_2_R) and in the heterodimer form A_2A_R-D_2_R. Previous we demonstrated that caffeine delays central fatigue through adenosine 2A receptors (A_2A_Rs) antagonism, striatal neuroplasticity and mitochondrial metabolism enhancement. In this study we demonstrate the participation of striatal A_2A_R-D_2_R in the ergogenic outcomes. Two-hundred swiss adult mice were treated systemically (i.p.) and centrally (striatum-st.) with caffeine, SCH58261 (A_2A_R antagonist), DPCPX (A_1_R antagonist) and haloperidol (D_2_R antagonist), and performed the open field test, treadmill incremental exercise and grip strength meter test We observed that A_2A_R-D_2_R antagonic effects are linked to motor control (*p* < 0.05) and enhancement of power (*p* < 0.05). However, the role of A_2A_R-D_2_R in the mechanism of strength generation seems to differ from motor control.

## Introduction

Adenosine is a modulator purine that acts in the central nervous system (CNS) and peripheral nervous system (PNS) [1]. There are four subtypes of adenosinergic receptors, the A_1_, A_2A_, A_2B_ and A_3_ (A_1_R, A_2A_R, A_2B_R e A_3_R), and they are coupled to G-proteins. The second messenger associated to adenosine-receptor binding is the activation or inhibition of adenylyl cyclase enzyme (AC) [2]. The A_1_R and A_2A_R have the highest affinity for adenosine [3].

Peripheral adenosinergic signaling participates in the blood supply through vasodilatation or angiogenesis [4], and decreased inflammatory response through suppression of cytokines production [5, 6]. Central adenosinergic signaling participates in the arousal state and fatigue [7]. The ergogenic effects of caffeine (1,3,7-trimethylxanthine) are due to central A_2A_Rs antagonism [7–9]. Recently, we demonstrated that striatum drives the ergogenic effects of caffeine (Alves et al., 2023).

The A_2A_Rs are highly expressed in the striatum [10], where they can be found co-localized with dopaminergic D_2_ receptors (D_2_R) in GABAergic medium spiny neurons (MSNs) [11]. The A_2A_Rs are coupled to stimulatory G_*s/olf*_ -proteins that activate the AC, increasing the production of AMPc (cyclic adenosine monophosphate) [12–14]. On the other hand, D_2_Rs are coupled to G_*i*_-proteins that inhibit the AC [15, 16].

Therefore, these receptors display antagonic effects on the same signaling cascade [17]. In addition to postsynaptic co-localization of A_2A_Rs and D_2_R, heterodimer forms of these receptors exist in the striatum [18, 19]. Since heterodimerization modifies the pharmacology of the receptor [20], it is important to investigate whether the heterodimer A_2AR_-D_2_R is involved in the ergogenic effects of caffeine and A_2A_R antagonism reported before.

Since the discovery of ergogenic effects of caffeine [7, 8, 21], selective A_2A_R and A_1_R antagonists were developed, such as SCH58261, and DPCPX, respectively. While SCH58261 has been linked to ergogenic effects [8, 22], studies relating DPCPX and fatigue remain scarce. Fatigue is defined as a subjective sense of tiredness and increased perceived effort compared to actual performance, or exhaustion [23, 24], and it can be measured objectively through fatigability. Fatigability is the magnitude or rate of change in a performance criterion relative to a reference value during a given task performance time or mechanical power measure [21, 25]. Therefore, the aim of this study is to investigate the peripheral and striatal effects of heterodimer A_2AR_-D_2_R antagonism in the fatigue, using caffeine, SCH58261, DPCPX and haloperidol (a D_2_R antagonist).

### Methods and experimental design

Two hundred male swiss mice (8 weeks old, 49.45 ± 1.5 g) were acquired from the Federal University of Santa Catarina (UFSC) and housed in collective cages (38 × 32 × 17 cm) with water and food *ad libitum*. Mice were maintained under a 12h light-dark cycle at a room temperature of 22 ± 1°C and monitored humidity. The experimental protocol was approved by the Animal Care and Use Committee of UFSC (IACUC-PP10519). Animals were randomly assigned to the experimental groups. To access fatigue, we evaluated behavior of mice in the open field test, grip strength meter test and treadmill incremental exercise.

### Drugs

For the systemic administration, all drugs were administered via intraperitoneal (i.p.) in a10 ml/kg volume. Caffeine (Sigma-Aldrich) and haloperidol (haldol®) were diluted in saline (NaCl 0.9%). SCH58261 (Sigma-Aldrich) and DPCPX (Sigma-Aldrich) were diluted in DMSO. Caffeine (6 mg/kg) were administered 15 min before tests, haloperidol (0.25 mg/kg) 30 min before, SCH58261 (1 mg/kg) 15 min, DPCPX (1 mg/kg) 30 min before. Saline and DMSO were administered in the respective control groups.

For the striatum administration, mice were anesthetized with ketamine/xylazine, and one cannulas was implanted in the right striatum (AP 0.5 mm; ML 2 mm and DV -3 mm) and another in the left striatum (AP 0.5 mm; ML -2 mm and DV -3 mm) (supplementary figure 1) for stereotaxy, according to Paxinos and Franklin’s mouse brain atlas [26]. Caffeine and haloperidol were diluted in saline (NaCl 0.9%). SCH58261 was diluted in DMSO. After one week of surgery, 4μL of caffeine (15μg) or saline (0.5μg) or SCH58261 (2μg) or DMSO (0.5μg) were injected into conscious animals using an infusion pump (2 μL/minute, Bonther®, Ribeirão Preto, Brazil) immediately before behavioral tests. Haloperidol (0.25 mg/kg) was administered 30 min i.p. before tests. Mortality rate was 30% (9 animals). All experimental doses were chosen based on pilot experiments and literature background [27–29]. Treadmill exercise could not be performed in animals after stereotaxic surgery because the length of cannulas were incompatible with the size of treadmill.

### Open field test

Motor control was evaluated in the open field apparatus (Insight® EP154). Mice were individually placed in the center of a circular apparatus (300×300mm) and allowed to explore for 5 min [30]. The number of crossings and rearings were manually analyzed.

### Grip strength meter test

Fatigue was evaluated in the grip strength meter apparatus and software (Bonther® 5 kgf). Each mouse was placed individually in the grip strength meter bar, and when both front paws were holding it steady, the experimenter gently pulled the tail in the other direction. It was performed in 4 trials with 10 seconds each. The final value is the average of the three best trials [31–34].

### Treadmill incremental exercise

Mice were habituated in a rodent treadmill for 3 days, and after 48h of rest, they performed the incremental test. Treadmill inclination (1.7 º) and shock (0.2 mA) were kept constant throughout the days. In the first day, animals performed 10 min of exercise in 5 meters/min (m/min), in the second day 5 min in 5 m/min + 5 min in 10m/min, and in the third day 10 min in 10 m/min intensity. After 48 h mice returned to the treadmill, and 5 m/min were add by the experimenter in each 3min of running, starting with 5 m/min. This incremental protocol was based in pilot experiments using measurement of blood lactate. Lactate was collected through submandibular vein puncture and analyzed using a lactimeter.

### Statistical analysis

All statistical analysis was performed in GraphPad Prism© (version 6.0.0 for Windows, GraphPad Software, San Diego, California, USA). One way ANOVA test (multiple comparisons) was used to investigate the differences among experimental groups. Outliers were detected and properly removed using GraphPad outlier calculator. p < 0.05 was considered statistically significant.

## Results

### The effects of systemic and central inhibition of A_2A_R-D_2_R on motor control

Caffeine (i.p.) increased number of crossings ([F_3, 36_] = 4.29, *p* < 0.05, fig. 1A) and rearings ([F_3, 36_] = 3.92, *p* < 0.05, fig. 1B) in the open field test. Haloperidol (i.p.) also increased number of crossings ([F_3, 36_] = 4.34, *p* < 0.05, fig. 1A) and rearings ([F_3, 36_] = 4.00, *p* < 0.05, fig. 1B) in the same test. However, the administration of caffeine plus haloperidol abolished statistical differences. Likewise, the administration of caffeine in the striatum (st.) of mice increased number of crossings ([F_2, 12_] = 9.67, *p* < 0.05, fig. 1C) and rearings ([F_2, 12_] = 6.08, *p* < 0.05, fig. 1D) in the open field test. Corroboring the systemic findings, the co-administration of caffeine (striatum) plus haloperidol (i.p.) decreased number of crossings ([F_2, 12_] = 9.67, *p* < 0.05, fig. 1C) and rearings ([F_2, 12_] = 7.79, *p* < 0.05, fig. 1D).

**Figure 1.**
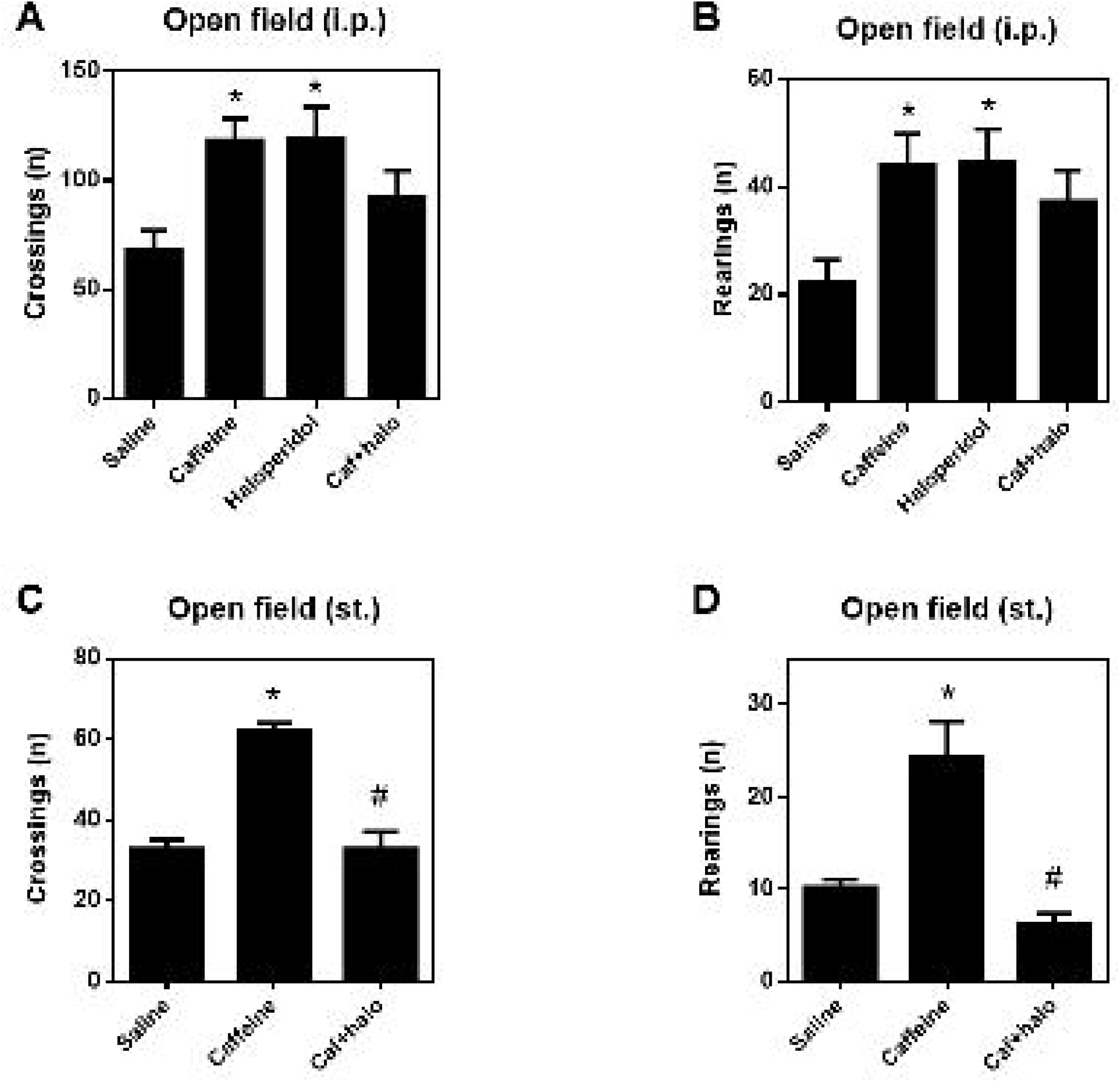
The effects of systemic and central administration of caffeine and haloperidol in the open field. Numbers of crosses (A) and numbers of rearings (B) after systemic administration. Numbers of crosses (C) and rearings (D) after central drug administration. * p < 0.05 compared to vehicle, # p < 0.05 compared to caffeine. One-way ANOVA with Tukey’s multiple comparison test, n=10 animals/group (systemic) and n=5 animals/group (central). i.p. = intraperitoneal; st.= stereotaxic; caff = caffeine; halo = haloperidol.

Similar to caffeine, systemic administration (i.p.) of SCH58261 increased number of crossings ([F_3, 36_] = 6.64, *p* < 0.05, fig. 2A) and rearings ([F_3, 36_] = 4.72, *p* < 0.05, fig. 2B) in the open field test, and the addition of haloperidol (i.p.) inhibited the psychostimulant effect of SCH58261 observed in the number of crossings ([F_3, 36_] = 4.20, *p* < 0.05, fig. 2A). As observed in the number of crossings ([F_3, 36_] = 7.59, *p* < 0.05, fig. 2B) and rearings ([F_3, 36_] = 4.44, *p* < 0.05, fig. 2B), systemic administration of DPCPX (i.p.) was not psychostimulant for mice, therefore it was not administered centrally. Striatal administration (st.) of SCH58261 also increased number of crossings ([F_2, 12_] = 11.45, *p* < 0.05, fig. 2C) and rearings ([F_2, 12_] = 4.07, *p* < 0.05, fig. 2D) in the open field test, and the co-administration of haloperidol (i.p.) inhibited these psychostimulant effect of SCH58261 in the number of crossings ([F_2, 12_] = 9.59, *p* < 0.05, fig. 2C) and rearings ([F_2, 12_] = 10.19, *p* < 0.05, fig. 2D).

**Figure 2.**
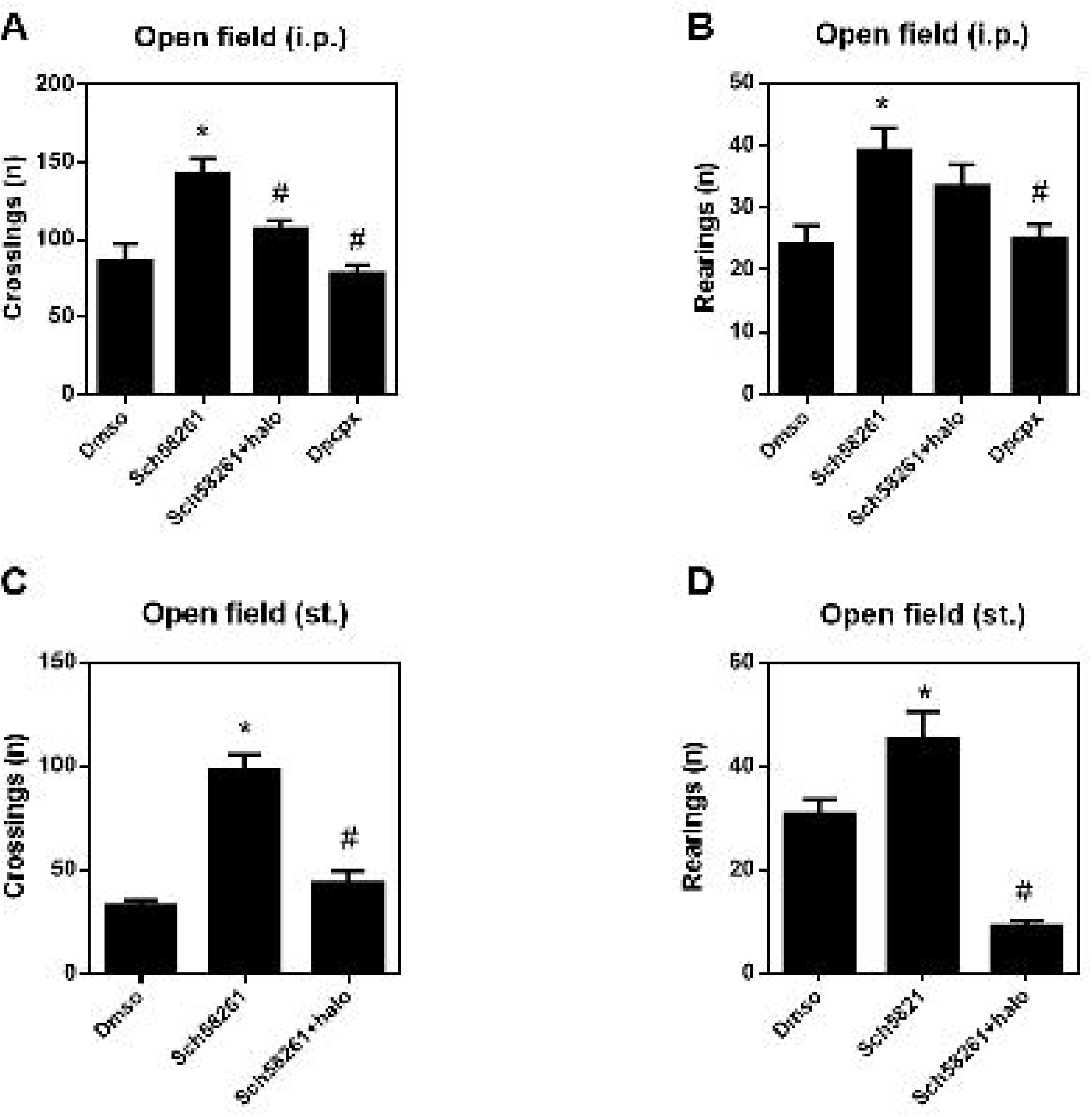
The effects of systemic and central administration of SCH58261 and haloperidol in the open field. Numbers of crosses (A) and numbers of rearings (B) after systemic administration. Numbers of crosses (C) and rearings (D) after central drug administration. * p < 0.05 compared to vehicle, # p < 0.05 compared to caffeine. One-way ANOVA with Tukey’s multiple comparison test, n=10 animals/group (systemic) and n=5 animals/group (central). i.p. = intraperitoneal; st.= stereotaxic; halo = haloperidol.

### The effects of systemic and central inhibition of A_2A_R-D_2_R on grip strength

Systemic administration (i.p.) of haloperidol per se ([F_3, 36_] = 5.75, *p* < 0.05, fig. 3A) and caffeine plus haloperidol ([F_3, 36_] = 5.11, *p* < 0.05, fig. 3A) decreased strength peak in the grip strength meter test in comparison to saline group. The co-administration of caffeine plus haloperidol also decreased peak strength in comparison to caffeine treatment ([F_3, 36_] = 7.78, *p* < 0.05, fig. 3A). Caffeine (i.p.) increased time of mice’ grip ([F_3, 36_] = 9.65, *p* < 0.05, fig. 3B), and haloperidol decreased this ergogenic effect of caffeine ([F_3, 36_] = 6.39, *p* < 0.05, fig. 3B), as corroborated by the area under curve of time x strength analysis ([F_3, 35_] = 11.49, *p* < 0.05, fig. 3C-D). Striatal administration (st.) of caffeine did not alter peak strength (fig 3E), but increased time of grip ([F_2, 27_] = 7.58, *p* < 0.05, fig. 3F) and area under curve ([F_2, 27_] = 6.63, *p* < 0.05, fig. 3G-H). However, unlike observed in the systemic administration, the co-administration of haloperidol (i.p.) also increased time ([F_2, 27_] = 6.64, *p* < 0.05, fig. 3F) and area under curve ([F_2, 27_] = 4.75, *p* < 0.05, fig. 3G-H) of mice.

**Figure 3.**
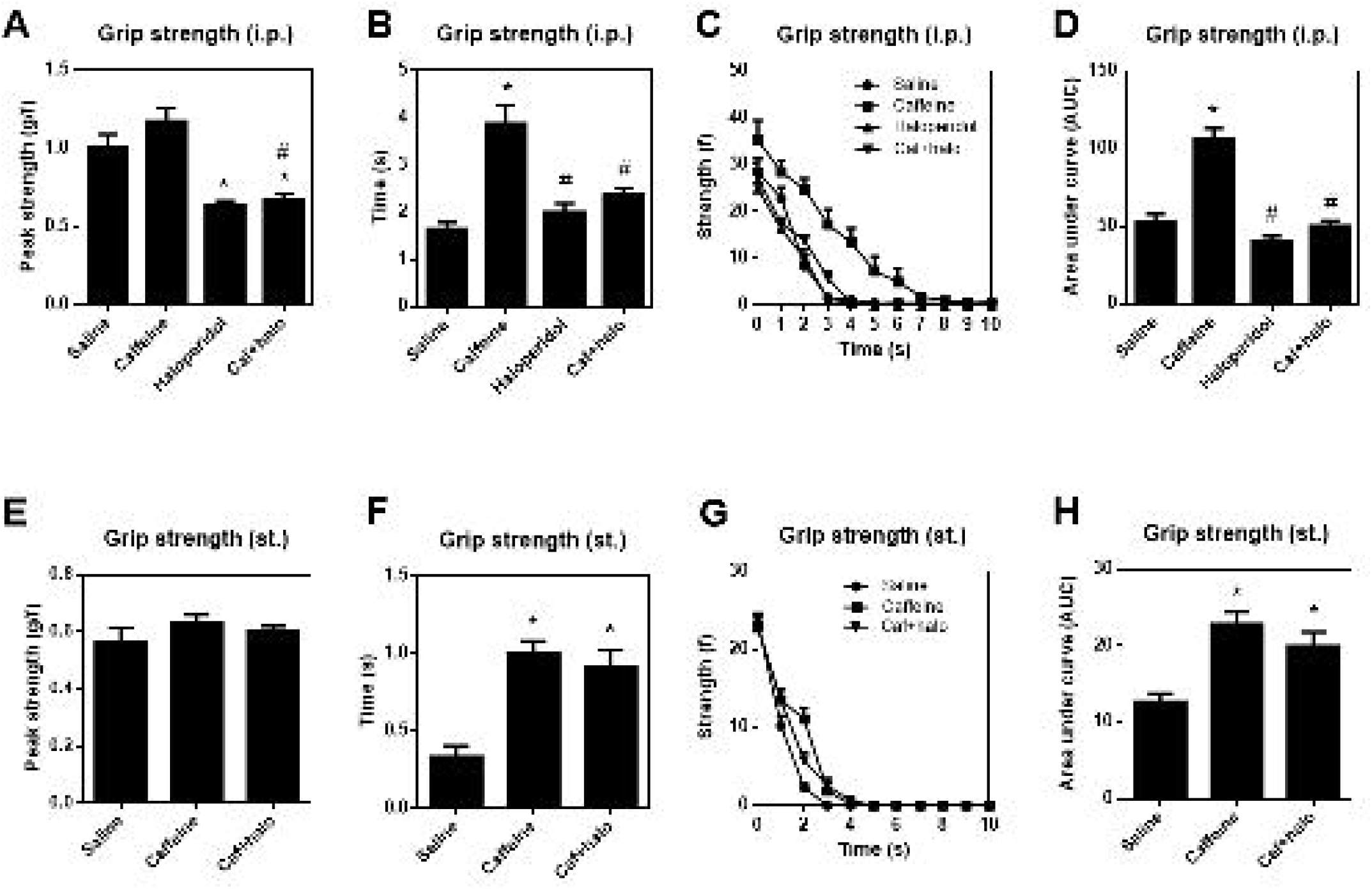
The effects of systemic and central administration of caffeine and haloperidol in the grip strength meter test. Relative peak force (A), grip time (B), time versus force curve (C) and area under the curve (D) of time versus force (C), after systemic drug administration. Relative peak force (E), hold time (F), time versus force curve (G) and area under the curve (H) of time versus force (G), after central drug administration. * p < 0.05 compared to vehicle, # p < 0.05 compared to caffeine. One-way ANOVA with Tukey’s multiple comparison test, n=10 animals/group (systemic) and n=5 animals/group (central). Caff = caffeine; halo = haloperidol.

SCH58261 (i.p.) did not alter peak strength (fig. 4A), but significantly increased time of grip ([F_3, 36_] = 4.80, *p* < 0.05, fig. 4B) in the grip strength meter test, and the area under curve of time x strength analysis ([F_3,36_] = 4.47, *p* < 0.05, fig. 4C-D). Once again the co-administration of haloperidol (i.p.) inhibited time of grip ([F_3, 36_] = 5.78, *p* < 0.05, fig. 4B), and the area under curve of time x strength analysis ([F_3,36_] = 5.64, *p* < 0.05, fig. 4C-D).

**Figure 4.**
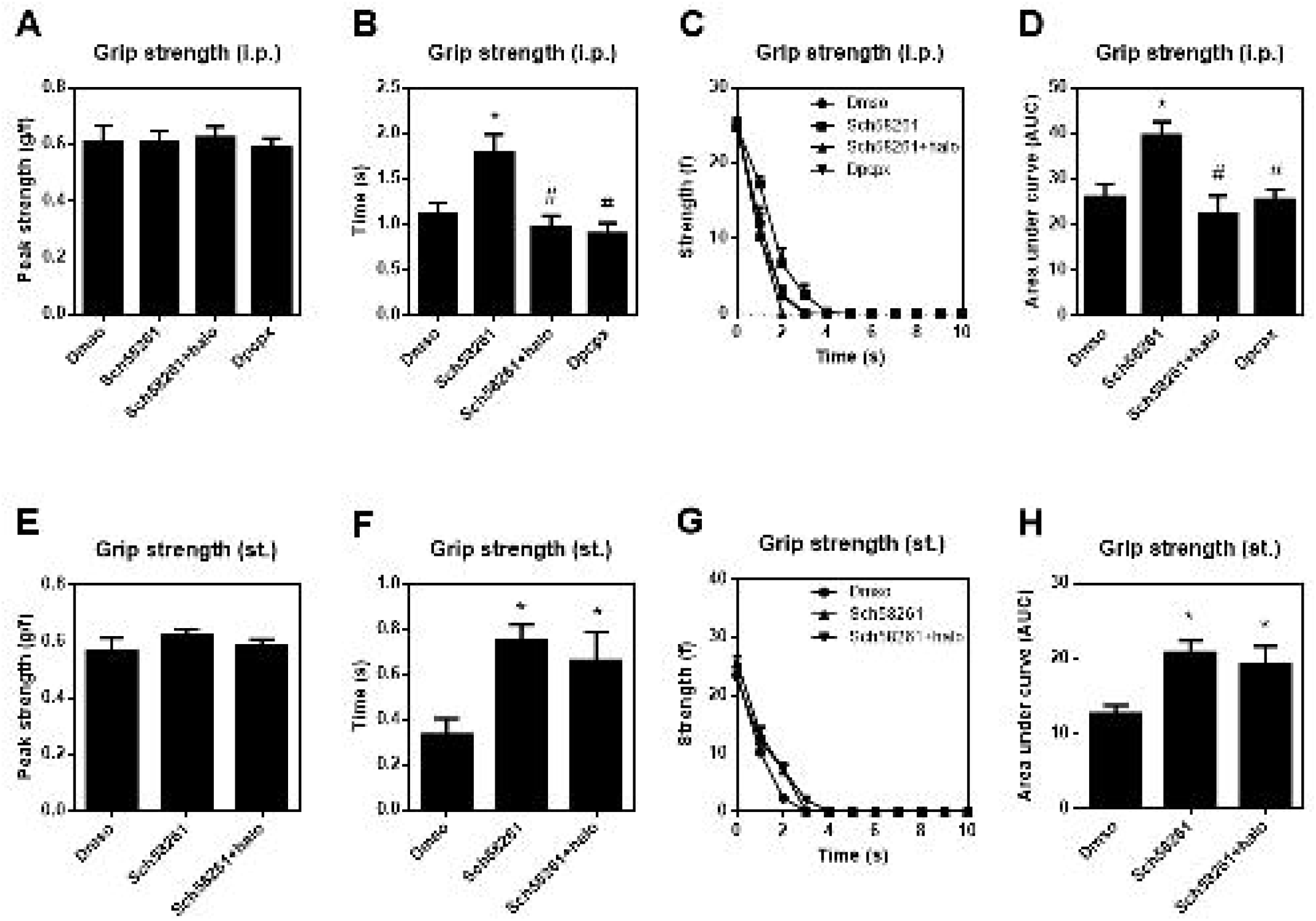
The effects of systemic and central administration of SCH58261 and haloperidol in the grip strength meter test. Relative peak force (A), grip time (B), time versus force curve (C) and area under the curve (D) of time versus force (C), after systemic drug administration. Relative peak force (E), hold time (F), time versus force curve (G) and area under the curve (H) of time versus force (G), after central drug administration. * p < 0.05 compared to vehicle, # p < 0.05 compared to caffeine. One-way ANOVA with Tukey’s multiple comparison test, n=10 animals/group (systemic) and n=5 animals/group (central). Caff = caffeine; halo = haloperidol.

Similar to striatal caffeine findings, central administration (st.) of SCH58261 increased time ([F_2,27_] = 4.46, *p* < 0.05, fig. 4F) and area under curve [F_2,27_] = 4.44, *p* < 0.05, fig. 4G-H), but also, the co-administration of haloperidol (i.p.) increased the same parameters ([F_2,27_] = 3.51, *p* < 0.05, fig. 4F) and [F_2,27_] = 3.53, *p* < 0.05, fig. 4G-H), respectively.

### The effects of systemic inhibition of A_2A_R-D_2_R on running exercise

Animal groups were not different in body weight (fig. 5A). Systemic caffeine (i.p.) increased blood lactate ([F_3,26_] = 4.05, *p* < 0.05, fig. 5B) and distance performed ([F_3,27_] = 4.11, *p* < 0.05, fig. 5D) on the test day. On the other hand, any treatment changed potency (fig. 5C and E). Co-administration of haloperidol (i.p.) abolished ergogenic effect of caffeine (fig. 5D).

**Figure 5.**
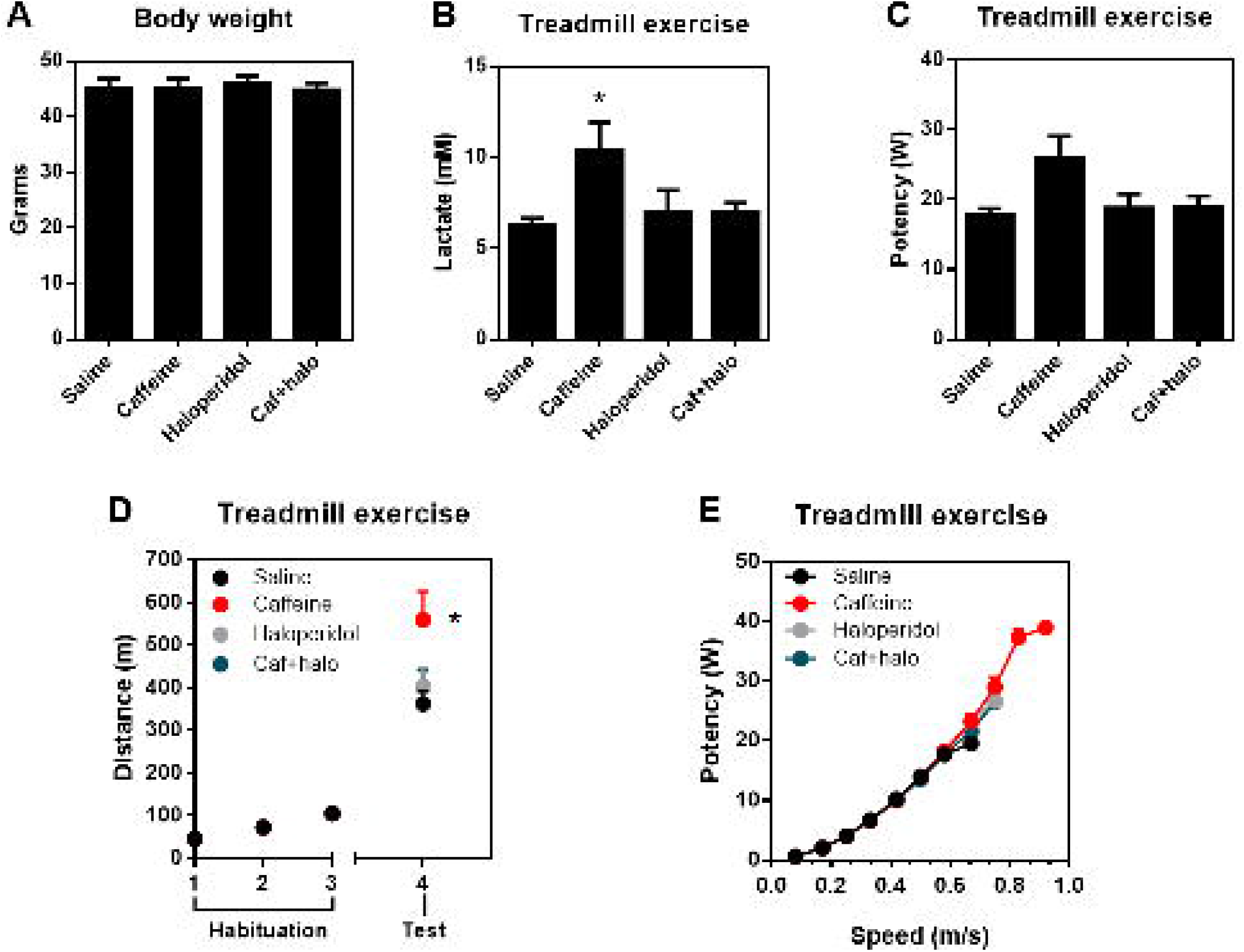
Effects of systemic administration of caffeine and haloperidol in the incremental treadmill test. Body weight (A), blood lactate (B), power (C), Distance performed (D) and power versus speed (E). * p < 0.05 compared to vehicle. One-way ANOVA with Tukey’s multiple comparison test, n=6-8 animals/group. Caff = caffeine; halo = haloperidol.

The secondary groups were not different in body weight (fig. 6A) and lactate level (fig. 6B). SCH58261 (i.p.) increased potency ([F_3,20_] = 5.33, *p* < 0.05, fig. 6C) and distance performed ([F_3,20_] = 6.74, *p* < 0.05, fig. 6D-E). The co-administration of haloperidol inhibited SCH58261 ergogenic effects in potency ([F_3,20_] = 5.61, *p* < 0.05, fig. 6C) and distance ([F_3,20_] = 5.00, *p* < 0.05, fig. 6D-E). In addition, DPCPX decreased mice potency ([F_3,20_] = 7.23, *p* < 0.05, fig. 6C) and distance peformed ([F_3,20_] = 6.51, *p* < 0.05, fig. 6D-E) compared to SCH58261 group.

**Figure 6.**
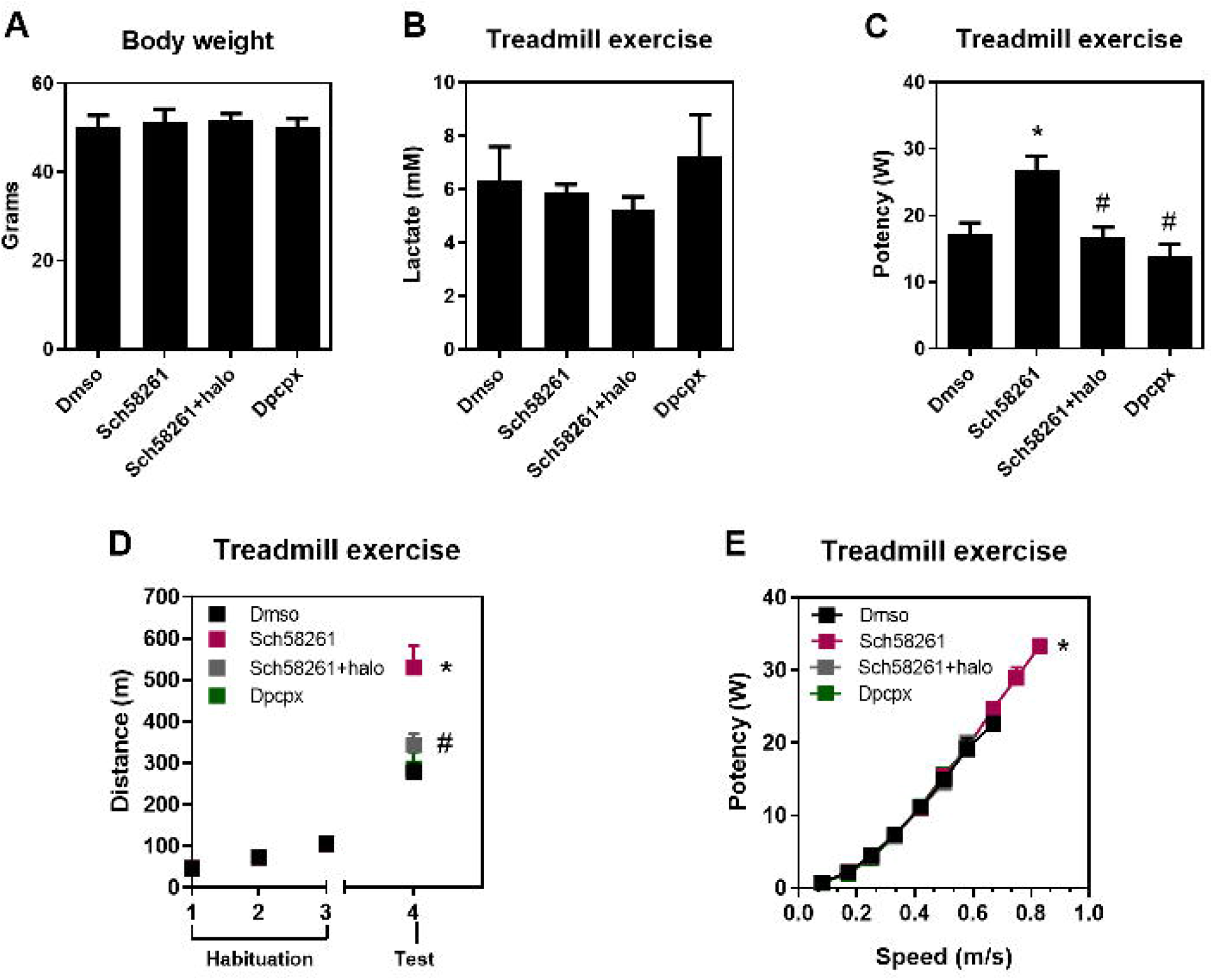
Effects of systemic administration of SCH58261 and haloperidol in the incremental treadmill test. Body weight (A), blood lactate (B), power (C), Distance performed (D) and power versus speed (E). * p < 0.05 compared to vehicle, # p < 0.05 compared to SCH58261. One-way ANOVA with Tukey’s multiple comparison test, n=6-8 animals/group. halo = haloperidol.

## Discussion

Our study strengthen that ergogenic and psychostimulant effects of caffeine are due to A_2A_R antagonism, specifically in the striatum, as confirmed by sistemic and local administration of caffeine and SCH 58261. Moreover, we demonstrated that A_1_R antagonism, through DPCPX, did not display any psychostimulant or ergogenic effect in mice. Curiously, heterodimer A_2A_R-D_2_R research revelead that mostly A_2A_R antagonism effects are due striatal D_2_R signaling cascade potentiation. However, we observed that dorsolateral striatal A_2A_R-D_2_R mediates motor control, but not grip strength, suggesting the involvement of other brain region in this mechanism.

Systemic administration of caffeine increased number of crossings and rearings in the open field test, and impulse magnitude in the grip strength meter test. These psychostimulant and ergogenic effects of caffeine corroborates the findings of our previous work [35]. In the current study, we extended the ergogenic effect of caffeine to treadmill exercise and added the participation of striatum. Other evidences support that caffeine increases 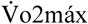, running power, and critical power in mice [8, 27]. The same psychostimulant and ergogenic effects were observed via striatal administration of caffeine. Psychostimulant and ergogenic effects of caffeine are linked to striatal neuroplasticity and enhancement of dopaminergic mitochondrial metabolism [8, 27]. Caffeine also increases presynaptic glutamate release and increases NMDA function in the striatum [13, 36], which may contribute to attenuation of fatigue during exercise.

Caffeine is a non-selective A_2A_R and A_1_R antagonist. Therefore, we investigated the effects of SCH 58261, and DPCPX on these parameters. Systemic administration of SCH58261 increased motor control of mice in the open field, impulse magnitude in the grip strength meter and running power in the treadmill exercise. Aguiar et al., 2020 [8], demonstrated that ergogenic effects of caffeine were abolished in global and forebrain A_2A_R knockout mice, showing that A_2A_R antagonism in forebrain neurons is responsible for the ergogenic action of caffeine [8]. On the other hand, systemic DPCPX was neither psychostimulant or ergogenic in mice, which is a novelty result. Striatal SCH58261 proved systemic results, showing that psychostimulant and ergogenic effects of A_2A_R antagonism are mediated in the striatum. The A_2A_Rs are located postsynaptically in striatopallidal GABAergic neurons, antagonizing D_2_R functions, and are also located presynaptically at corticostriatal terminals, facilitating glutamate release. Corroborating our findings [35], Shen et al., 2013, reported that A_2A_Rs in the glutamatergic terminals regulate psychostimulant actions of drugs through the integration of striatal GABAergic, dopaminergic and glutamatergic signaling [37].

Approaching the heterodimer A_2A_R-D_2_R antagonic effects, we found that psychostimulant and ergogenic effects of A_2A_R antagonism in the open field and treadmill exercise performance are due striatal D_2_R signaling cascade potentiation. Salamone et al., 2012, investigated the role of heterodimer A_2A_R-D_2_R in effort-related processes and observed that in addition to dopamine, particularly in the nucleus accumbens, adenosine through activation of A_2A_R, also participates in regulating these processes [38]. Besides the role of A_2A_Rs in the integration of neurotransmitters signaling stated before [37], the opposing actions over AC signaling cascade, the A_2A_Rs also facilitate neuronal activity in the striatum by improving dopaminergic signaling, through increased D_2_R availability and enhancement of dopamine neurotransmission [39]. However, while systemic and striatal antagonism of A_2A_R improved impulse magnitude in the grip strength meter test, the D_2_R antagonism (by haloperidol) only abolished this effect via systemic, but not when administered via striatum. Our study explored the dorsolateral striatum, this last finding may suggest that D_2_R from other regions, for example nucleus accumbens, in the ventral striatum, participates in the strength generation. Thus, future studies should be performed to elucidate this mechanism.

## Conclusion

We demonstrated that striatal heterodimers A_2A_R-D_2_Rs are linked to psychostimulant and ergogenic effects of adenosine antagonism in mice.

## Declarations Ethics approval

Experimental protocol was approved by the Animal Care and Use Committee of Federal University of Santa Catarina -UFSC (IACUC-PP10519).

## Consent for publication

The authors have agreed to publish this manuscript.

## Avaliability of data and materials

Data is available at OSF (DOI 10.17605/OSF.IO/E5DX8; DOI 10.17605/OSF.IO/69ZB5 and DOI 10.17605/OSF.IO/9QDKW).

## Authors’ contributions

Alves, A.C.B. performed the experiments, wrote the manuscript main text and prepared the figures. Santos, N.S., Tavares, A.P., and Panatta, G. assisted in the experiments. Aguiar, A.S.Jr. designed the methodology, revised the text, figures and statistical analysis.

## Acknowledgements

We thank the funding agencies and the LaBioEx co-workers.

## Author information

### Meet the author guidelines

Ana Cristina de Bem Alves graduated with BSc in physical therapy from the Federal University of Santa Catarina (UFSC), with an internship at the University of Northern Iowa and the University of Pittsburgh. She has a master’s degree in neurosciences and is currently finishing her Ph.D. at UFSC. She has expertise in neurodegenerative diseases, emphasizing neuroinflammation and the adenosinergic system. In the past two years, her studies have been directed to the potential of adenosinergic A2A receptors (A2AR) antagonism in managing fatigue. She is dedicated to finding new therapeutic options for neurological patients, and improving their quality of life.

---

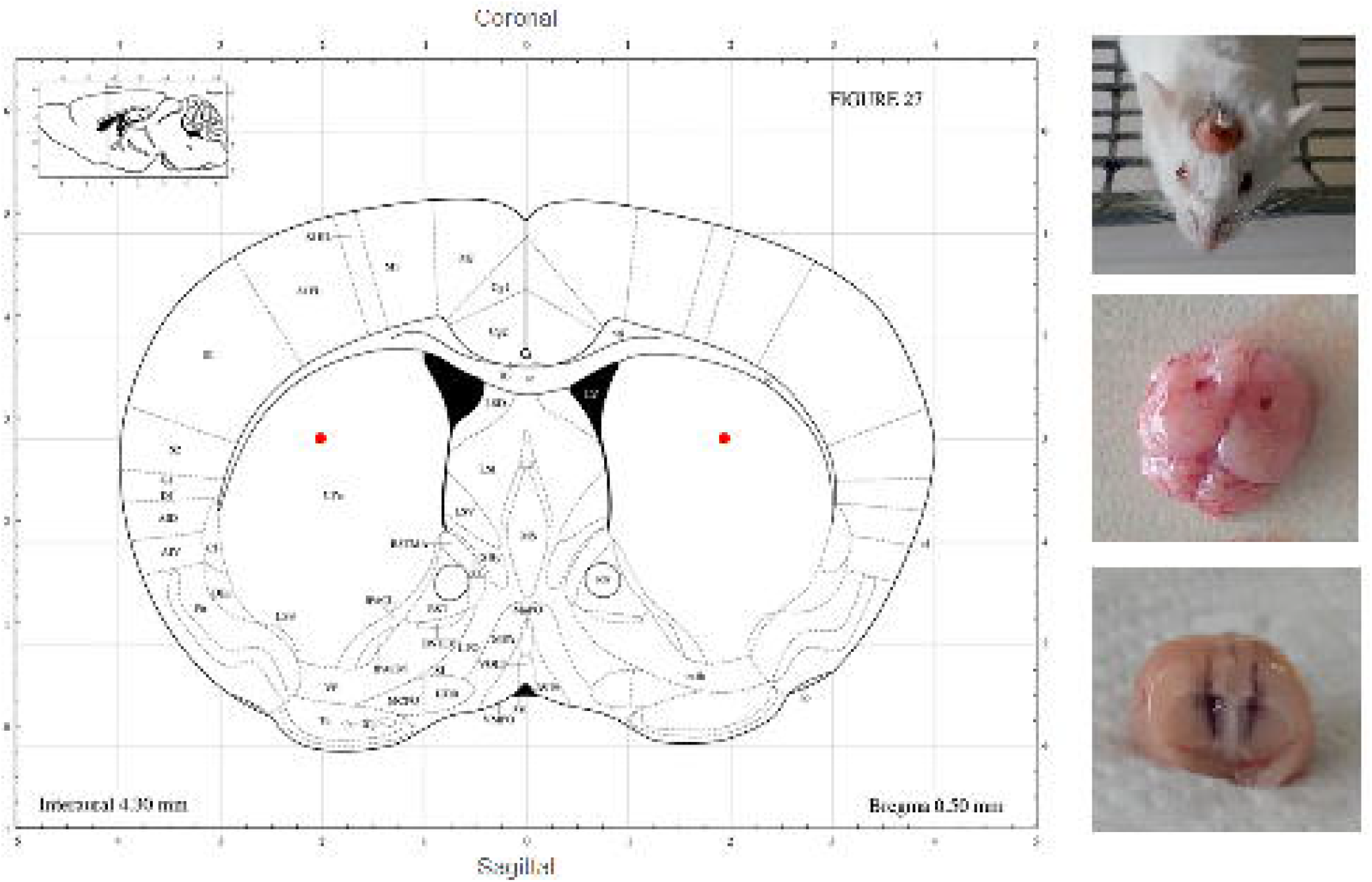

